# Loss of transcriptional heterogeneity in aged human muscle stem cells

**DOI:** 10.1101/2022.05.11.491465

**Authors:** Emilie Barruet, Katharine Striedinger, Pauline Marangoni, Jason H. Pomerantz

## Abstract

Age-related loss of muscle mass and function negatively impacts healthspan and lifespan. Satellite cells function as muscle stem cells in muscle maintenance and regeneration by self-renewal, activation, proliferation and differentiation. These processes are perturbed in aging at the stem cell population level, contributing to muscle loss. However, how representation of subpopulations within the human satellite cell pool change during aging remains poorly understood. We previously reported a comprehensive baseline of human satellite cell (Hu-MuSCs) transcriptional activity in muscle homeostasis describing functional heterogenous human satellite cell subpopulations such as CAV1+ Hu-MUSCs (Barruet et al., 2020). Here, we sequenced additional satellite cells from new healthy donors and performed extended transcriptomic analyses with regard to aging. We found an age-related loss of global transcriptomic heterogeneity and identified new markers (*CAV1, CXCL14, GPX3*) along with previously described ones (*FN1, ITGB1, SPRY1*) that are altered during aging in human satellite cells. These findings describe new transcriptomic changes that occur during aging in human satellite cells and provide a foundation for understanding functional impact.

## INTRODUCTION

Aging in skeletal muscle is characterized by a decline in muscle mass and regenerative capacity manifested in humans by decreased muscle strength and slow healing after injury (Curtis, Litwic, Cooper, & Dennison, 2015; Distefano & Goodpaster, 2018). At the cellular level, muscle fiber growth, turnover and regeneration are driven by muscle stem cells also known as satellite cells (Chang & Rudnicki, 2014), characterized by the expression of PAX7. Satellite cells undergo activation and proliferation upon injury, and then either differentiate to generate new muscle fibers or return to a quiescent state to reconstitute the satellite cell pool. Therefore alterations in satellite cells during aging could underlie associated changes in muscle bulk and function.

There has been discordance in the literature with several studies reporting age-related loss in number and function of satellite cells (Chakkalakal, Jones, Basson, & Brack, 2012; Conboy, Conboy, Smythe, & Rando, 2003; Sousa-Victor et al., 2014) while others have reported no significant reduction or change (Arpke et al., 2021; Schäfer, Zweyer, Knauf, Mundegar, & Wernig, 2005). We observed a modest decrease in satellite cell number in samples from elderly (>81 years) human individuals (Garcia et al., 2018). Age can also cause intrinsic changes of satellites cells, their niche or both (Brack & Muñoz-Cánoves, 2016; Chakkalakal et al., 2012; Ermolaeva, Neri, Ori, & Rudolph, 2018; García-Prat et al., 2016; Lukjanenko et al., 2016; Rozo, Li, & Fan, 2016). Recent advances in single-cell genomics have allowed the discovery of novel aspects of aging in different tissues, which includes changes in cell heterogeneity, distribution of cellular states and gene expression levels (Angelidis et al., 2019; Consortium, 2020; Jacob C Kimmel et al., 2019; Kowalczyk et al., 2015). Studies in mice profiling the transcriptome of muscle stem cells along differentiation pathways have revealed age-related changes such as a decrease in expression of extracellular matrix (ECM), migration and adhesion genes (Jacob C. Kimmel, Yi, Roy, Hendrickson, & Kelley, 2021). Most prior intrinsic satellite cell aging studies have been performed in mice, with few efforts to translate those findings to humans (Snijders et al., 2015). Thus, a comprehensive characterization of the impact of aging on human satellite cells is still lacking.

We previously demonstrated that under homeostatic conditions, human satellite cells are transcriptionally heterogeneous, which enabled us to separate functionally distinct human satellite cell subpopulations (Barruet et al., 2020). Therefore, a comprehensive understanding of how the repartition of these subpopulations and their gene expression vary along with aging is now feasible._In this study, we used single cell RNA sequencing of satellite cells from additional human muscle samples to further analyze existing datasets with regard to aging. We demonstrate that there is a loss of transcriptional heterogeneity with aging and identify new genes that are differentially expressed during aging.

## RESULTS

### 1 Decreased transcriptional heterogeneity in Hu-MuSCs during aging

Our previous work identified functionally heterogenous human satellite cell subpopulations. We asked whether the distribution of those subpopulations or their transcriptome are altered in aging. We performed single cell RNA sequencing of highly purified human satellite cells from new healthy donors (**Figure 1-Source Data 1**) and pooled them with our previous dataset (Barruet et al., 2020). Our analysis workflow is described in **Figure 1A**. The single cell sequencing data for each sample were analyzed using SCANPY (F. Alexander Wolf, Angerer, & Theis, 2018), merged into their respective age group using the BBKNN (batch balanced k nearest neighbors) integration algorithm (Polański et al., 2019) to remove batch effect, and visualized in uniform manifold approximation and projection (UMAP) graphs (**Figure 1-figure supplement 1A-B**). Samples were distributed in 3 age groups: young [<30 y.o.], adult [35-66 y.o.] and aged [>70 y.o.]. All samples within each group were pooled. We identified 12, 9 and 10 clusters for the young, adult and aged groups, respectively. Myogenic, cycling and stemness genes were expressed as previously described (Barruet et al., 2020) in each age group. Moreover, similar cluster markers such as AP-1 transcription factor unit (*JUN, FOS*), *COL1A1* (Collagen Type Alpha 1 Chain), *SOX8* (Sry-Box Transcription Factor 8), *IGFBP7* (Insulin Like Growth Factor Binding Protein 7), *MX1, HSPA1A* (Heat Shock Protein A, Hsp70)) or *CAV1* (Calveolin-1) were found in each age group (**Figure 1-figure supplement 1B**). To compare each age group to another, we used INGEST (De Santis, Etoc, Rosado-Olivieri, & Brivanlou, 2021; Stuart et al., 2019). Unlike BBKNN or CCA (Canonical Correlation Analysis, e.g. in Seurat) where datasets are integrated in a symmetric way, INGEST integrates asymmetric datasets into a ‘reference’ annotated dataset. We found that the clusters obtained in our grouped young samples were robustly defined by accepted markers, in addition to having the greatest transcriptional variability (estimated through a higher number of clusters). Hence, we used this young group as our ‘reference’ annotated dataset to best identify potential differences in the transcriptional signatures induced by aging (**Figure 1-figure supplement 1C**). This approach allowed us to detect the biological variation observed with aging.

**Figure 1:**
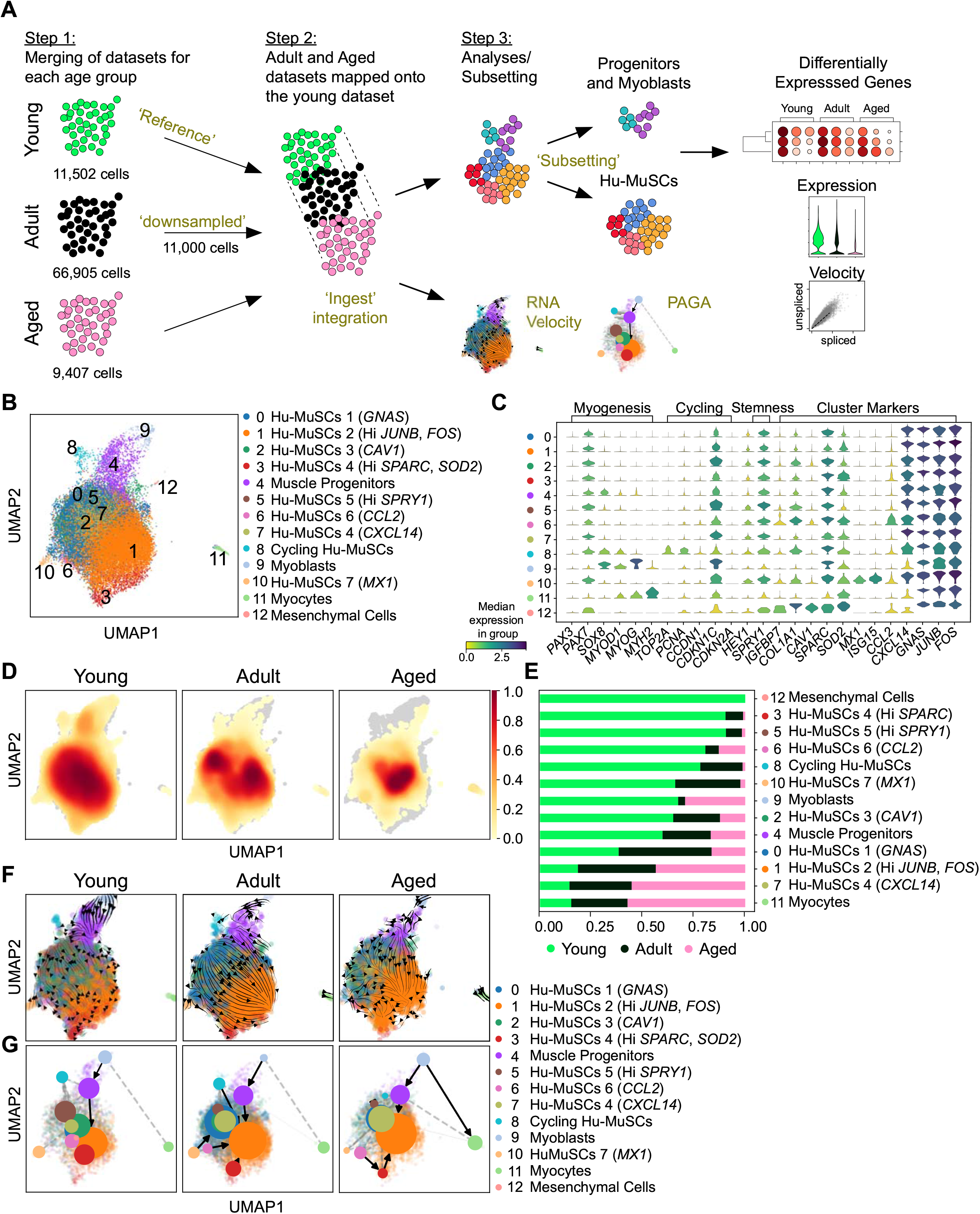
Distribution of Hu-MuSC subpopulations during adult aging. (A) Workflow of the analysis. (B) UMAP of merged age groups using INGEST with labeled clusters. (C) Violin plots displaying the expression of myogenic, cycling, stemness and cluster marker genes for each cluster. (D) UMAP density plot for each age group (Young, Adult, Aged). (E) Proportion plot of cells assigned to each age group according to each cluster. (F) RNA velocities projected onto the UMAP clusters for each age group. (G) PAGA analysis for the different age group. Weighted edges correspond to the connectivity between two clusters.

Prior to mapping, the adult group of cells was downsampled to 11,000 cells to remove co-founder effect resulting from differences in cell number among the 3 age groups. Among the 12 clusters, clusters 0-3, 5-7 and 10 consisted of quiescent Hu-MuSCs while cluster 4, 8, 9, 11 and 12 consisted of muscle progenitors, cycling Hu-MuSCs, myoblasts, myocytes and mesenchymal cells, respectively. We confirmed that each Hu-MuSC cluster was found to have a unique transcriptomic fingerprint. Cluster 0 was characterized by the upregulation of *GNAS* and cluster 1 by the upregulation of *JUNB* and *FOS*. Cluster 2 contained the *CAV1* expression cells while cells expressing high levels of *SPRY1* (Sprouty RTK Signaling Antagonist 1) were found in cluster 5. Cluster 6 and 7 consisted of cells expressing cytokines *CCL2* and *CXCL14*. Finally, the recently described (Scaramozza et al., 2019) *MX1* satellite cell subpopulation was identified as cluster 10 (**Figure 1B-C**).

The INGEST integration allowed us to compare the distribution of the young, adult and aged Hu-MuSCs among the different clusters. A UMAP based-density plot revealed a decrease of cluster coverage with aging (**Figure 1E** and **Figure 1-figure supplement 1D**). The majority of aged Hu-MuSCs were located in cluster 1 (65%, Hi *JUNB, FOS*) and cluster 7 (10.9%, *CXCL14*) while young Hu-MuSCs were distributed among a larger number of clusters such as cluster 2 (14.8%, *CAV1*), cluster 3 (13.3%, Hi SPARC, (Secreted Protein Acidic and Cysteine Rich, an ECM protein (Jørgensen et al., 2017)), SOD2 (Superoxide Dismutase 2)), cluster 5 (3.9% Hi *SPRY1*) and cluster 6 (2.4%, *CCL2*). Adult cells were predominantly present in cluster 0 (33.5%), 1 (48.9%) and 7 (5.1%) (**Figure 1D-E** and **Figure 1-figure supplement 1D-E**). Thus, distribution of cells per cluster varies with aging and there is a relative loss of transcriptional heterogeneity in aged Hu-MuSCs.

Since the distribution of cells per cluster appears affected by aging, we aimed to investigate the direction and speed of movement of cells in clusters inferred by RNA velocities (La Manno et al., 2018). To understand the cellular dynamics of Hu-MuSCs and population kinetics during aging, we applied the scVelo and partition-based graph abstraction (PAGA) trajectory algorithm (Bergen, Lange, Peidli, Wolf, & Theis, 2020). scVelo and PAGA analyses suggest that in young cells, cluster directionality is heterogenous, contrary to adult and aged cells where cell states appear to commit to cluster 1 (**Figure 1F,G**). Since the weighted edges correspond to the connectivity between two clusters in the PAGA analysis, we were able to more closely investigate the connectivity between clusters. In the three age groups we found a strong connection axis between clusters 1 (Hu-MuSCs, hi *JUNB, FOS*), 4 (Muscle Progenitors) and 9 (Myoblasts) with directional kinetics toward the less differentiated states. Aged cells from other Hu-MuSC clusters (0, 2, 3, 6, 10) converged toward cluster 1 while in the young age group, Hu-MuSC clusters appear to be in a non-convergent steady state. We also found that cells from cluster 7 (*CXCL14*) in the young group were not connected to any other cluster in contrast to adult and aged cells (**Figure 1G**). These findings suggest that aging Hu-MuSCs have a decrease in transcriptional hetero-geneity characterized by cluster specificity and convergent RNA velocity.

### 2 Extracellular-matrix and adhesion gene expression decreases with aging in Hu-MuSCs

Since we observed a loss in transcriptional heterogeneity, we asked which genes may be differentially expressed during aging. We excluded activated, differentiated and non-myogenic cells and focused on the Hu-MuSC clusters (0-3, 5-7 and 10) solely to assess modulations in transcriptional signatures and highlight age-related modifications (**Figure 2A,B**). We then identified differentially expressed genes for the three age groups, where each age group was compared to one another. Notably we found collagen genes (i.e. *COL3A1*) to be significantly expressed in young Hu-MuSCs, while *DUSP1* (can inactivate MAPK proteins and has been reported to increase upon Hu-MuSC activation in culture (Charville et al., 2015)) and *ZFAND5* (a proteosome activator (Lee, Takayama, & Goldberg, 2018)) were significantly expressed in adult Hu-MuSCs, and *CXCL14* and *GPX3* (Glutathione Peroxidase 3, a retinoid-responsive gene (El Haddad et al., 2012)) in aged Hu-MuSCs (**Figure 2C**). These results also corroborate previously described genes affected by aging such as *FN1* (Fibronectin 1), *ITGB1* (Integrin Subunit Beta-1), *SPRY1* (Lukjanenko et al., 2016; Rozo et al., 2016; Shea et al., 2010) which were decreased with aging. Moreover, expression of other ECM and adhesion related genes (*CAV1*, *COL1A2*) decreased aged cells. The expression of mTor pathway target genes such as *BCL2* and *VEGFA* (Vascular Endothelial Growth Factor A) was increased in adult and aged Hu-MuSCs (**Figure 2D**).

**Figure 2:**
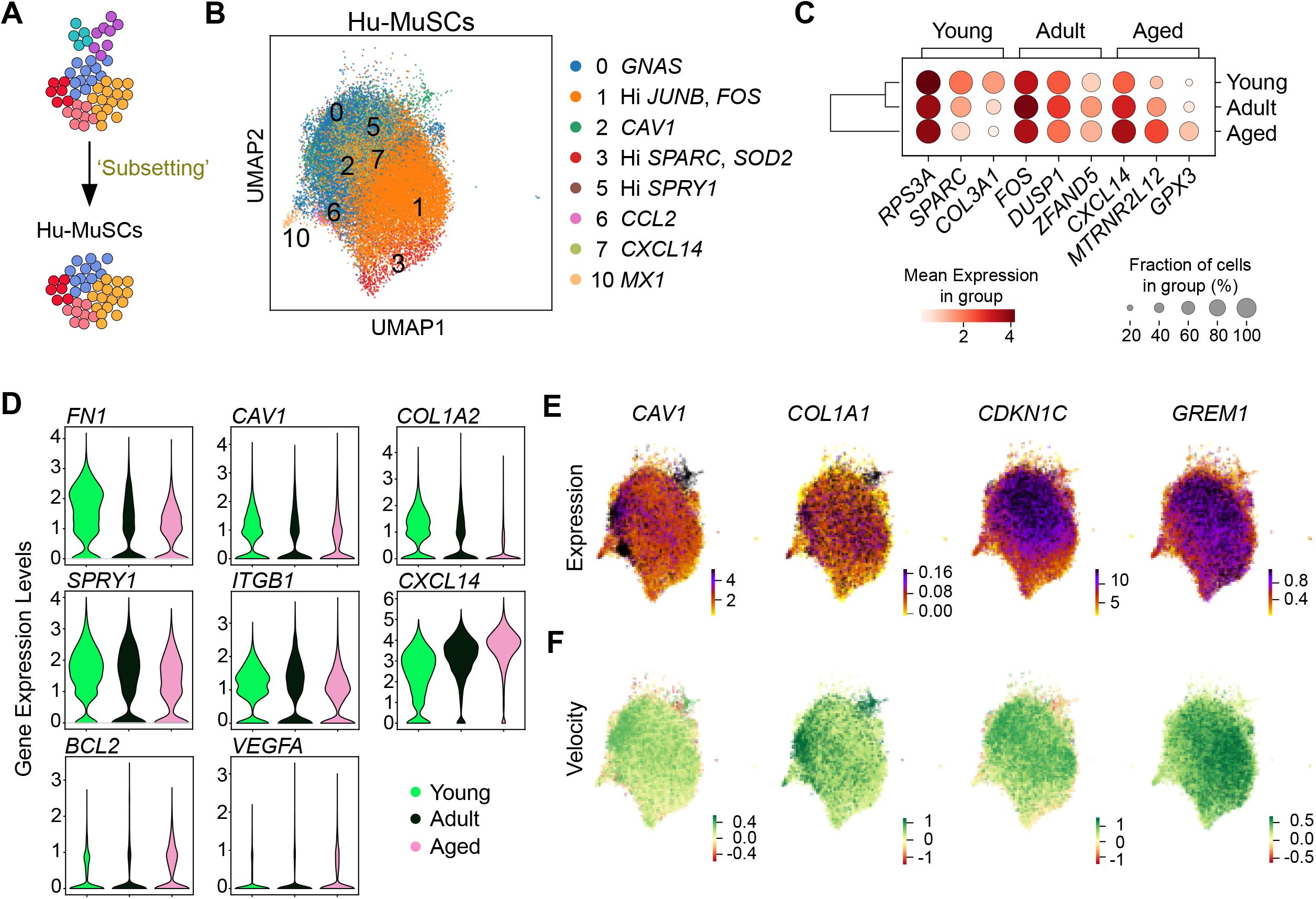
Gene expression and velocity analyses of Hu-MuSCs of young, adult and aged groups. (A) Schematic of the sub-clustering. Only Hu-MuSCs were analyzed. (B) UMAP of merged age groups for only the human muscle stem cells (cluster 0-3, 5-7 and 10). (C) Dot plot displaying the top 3 genes differentially expressed for each age group in the whole Hu-MuSCs subset. (D) Violin plots of the expression of genes altered with aging. (E) Expression of relevant genes that inferred age-specific differential velocity. (F) Velocity of genes from (E).

Since the distribution of cells among the different clusters as well as population kinetics changed with aging, we asked if the inferred directions described in **Figure 1F** were supported by any specific genes or transcriptional program. In comparing the three age groups, we found that genes including *CAV1*, *COL1A1*, *CDKN1C* (Cyclin Dependent Kinase Inhibitor 1C, regulates cell proliferation (Mademtzoglou et al., 2018)) and *GREM1* (Gremlin1, a BMP antagonist (Borok, Mademtzoglou, & Relaix, 2020)) had age-specific differential velocity expression meaning that those genes were transcribed at significantly higher or lower levels compared to their age group counterpart (**Figure 2E,F**). The UMAP of velocity expression showed that *CAV1* and *COL1A1* had increased velocity in the clusters enriched in the young samples (cluster 2, 3 and 5) while *CDKN1C* and *GREM1* displayed increased velocity in clusters enriched with the adult (cluster 0) and aged (cluster 1) samples, respectively (**Figure 2F**). We also found additional genes with an age-specific significant differential velocity. These include *DIO2* (Type 2 Iodothy-ronine Deidinase, coverts thyroid prohormone (Buroker, 2014)), *EDN3* (Endothelin-3, mediates the release of vasodilators (Kawanabe & Nauli, 2011)), *NPTX2* (Neuronal Pentraxin 2, which affect tumor progression (Z. Wang et al., 2020)), *KLF6* (Krueppel-like Factor 6, a tumor suppressor (Tetreault, Yang, & Katz, 2013)), *LPL* (Lipoprotein Lipase, involved in lip metabolism (Wu, Kersten, & Qi, 2021)) and *MAP1B* (Microtubule Associated Protein (Halpain & Dehmelt, 2006)). *DIO2, LPL, MAP1B, EDN3* were top ranked genes that explained the resulting vector field of young Hu-MuSCs with an increase in velocity. *NPTX2* and *KLF6* velocity were increased in cluster 1 and 7, clusters associated with adult and aged Hu-MuSCs (**Figure 2-figure supplement 1** and **Figure 2-Source Data 1**).

The gene ontology (GO) term analysis of differentially expressed genes in young, adult and aged Hu-MuSCs revealed an enrichment in ECM terms in young Hu-MuSCs although they were decreased in aged Hu-MuSCs. Type I / IFNy signaling were decreased in young Hu-MuSCs. Terms associated with muscle processes were increased in aged Hu-MuSCs and downregulated in adult Hu-MuSCs. An enrichment of cellular response to oxygen levels and hypoxia terms was also detected in aged Hu-MuSCs (**Figure 3**).

**Figure 3:**
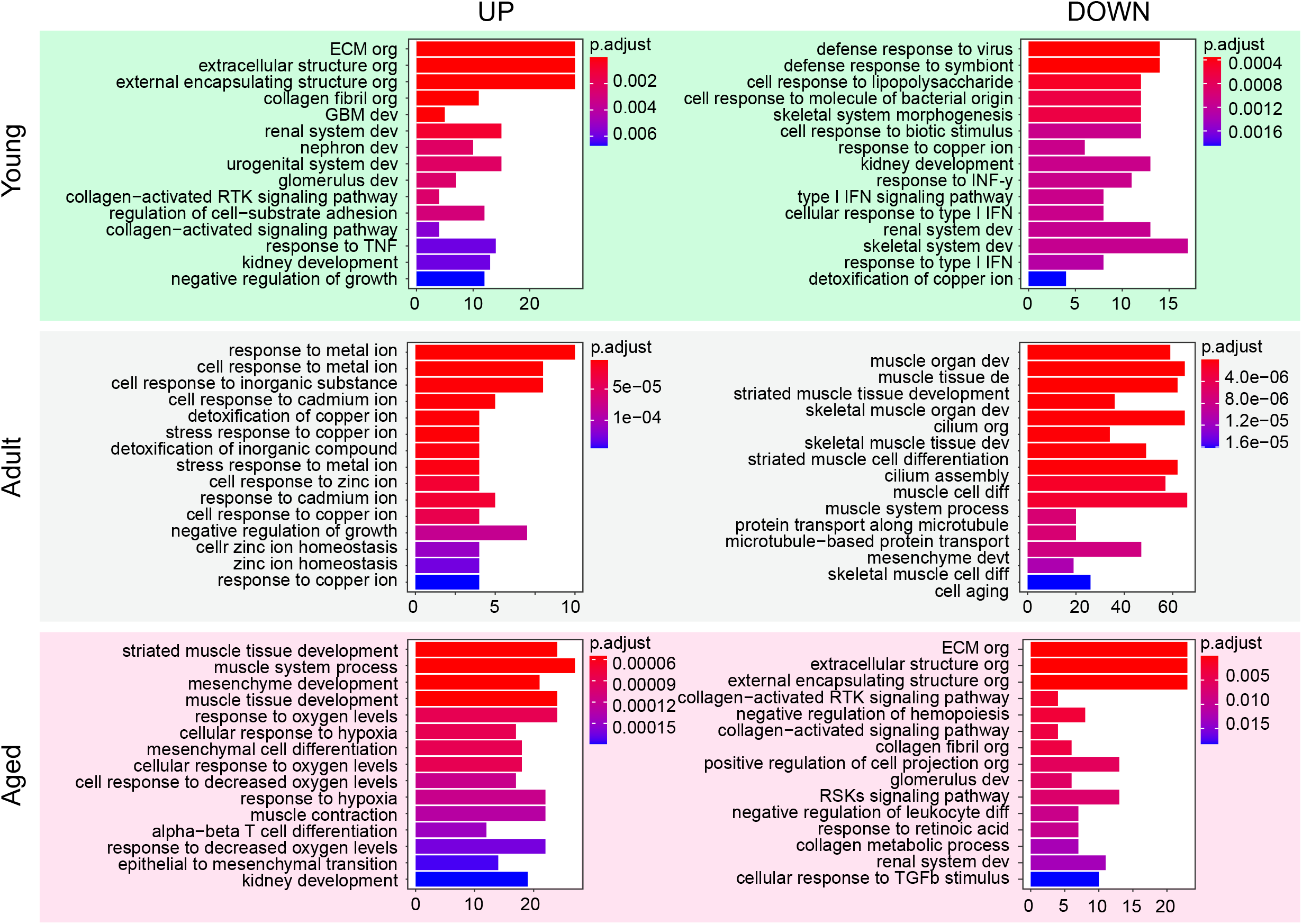
Gene ontology enrichment upon aging in Hu-MuSCs. Bar plots of gene ontology analysis of differentially up- and down-regulated genes in Hu-MuSCs for each age group.

To summarize, our approach resulted in the identification of novel aging-related markers such as *CAV1* and *GREM1* while verifying in Hu-MuSCs the expression levels of genes that have been previously associated with murine aging alterations.

### 3 scRNAseq analysis also reveals transcriptomic changes during aging in muscle progenitor cells, cycling Hu-MuSCs and myoblasts

Since our dataset also contained a large fraction of differentiated cells, we separately analyzed clusters encompassing the cycling Hu-MuSCs (cluster 8), muscle progenitors (cluster 4) and my-oblasts (cluster 9) (**Figure 4A**). Top differentially expressed genes in the muscle progenitors included *MEST* (Mesoderm Specific Transcript, a negative regulator of adipocyte differentiation (Karbiener et al., 2015)), *HSPG2* (Heparan Sulfate Proteoglycan 2, encodes for secreted molecule perlecan, deposited on all basement membrane (Martinez, Dhawan, & Farach-Carson, 2018)), *OLFM2B* (Olfactomedin 2b, a regulator for TGF-b (Shi, Guo, & Chen, 2014)) and *SPARC* for the young samples; *FOS*, Metallothionein genes (*MT1E* and *MT2A*) and *SPARCL1* (SPARC-like protein 1, an ECM protein that has been described as a differentiation promotor of C2C12 cells (Y. Wang, Liu, Yan, Li, & Tong, 2019)) for the adult samples; and *NUPR1* (Nuclear Protein 1 Transcription Regulator, a repressor of ferroptosis (J. Liu et al., 2021)), *TRDN* (Triadin, plays a role in muscle excitation-contraction (Marty & Fauré, 2016)), *MTRNR2L12* (an isoform of humanin (Bik-Multanowski, Pietrzyk, & Midro, 2015)) for the aged samples. As with our Hu-MuSC explorations, GO term analysis of differentially expressed genes for differentiated cell clusters showed an enrichment of ECM and cell matrix adhesion terms in young muscle progenitors, along with an enrichment of muscle cell differentiation and interferon gamma terms in aged muscle progenitor cells. Similar analysis for the cycling Hu-MuSCs revealed a significant increase of *TUBA1B* (Tubulin Apla-1B, a cytoskeleton protein (Q. Q. Xu et al., 2020)), *TYMS* (Thmidylate Synthetase, a critical enzyme for DNA replication and DNA repair (Varghese et al., 2019)), *H2AFV* (H2A.Z Variant Histone 2), *STMN1* (Stathmin1, a microtubule-binding protein (Jun Liu et al., 2021)) transcripts levels in young cells, similar to that of *SPARCL1, CXCL14, ZFP36* (ZFP36 Ring Finger Protein), *EIF1* (Eukaryote Translation Initiation Factor 1) in adult cells and Proteosome proteins (*PSMB10* and *PSMB9), PRDX1* (Peroxiredoxin 1, an antioxidant enzyme (Ding, Fan, & Wu, 2017)) and *S100A16* (S100 Calcium Binding Protein A16) in aged cells. GO term analysis showed enrichment in DNA replication and cell cycle terms in young cycling Hu-MuSCs, cellular response to metal ion terms in adult cells and enrichment in antigen processing and Wnt signaling pathway in aged cycling Hu-MuSCs (**figure 4-figure supplement 1**). Finally, *CDKN1C, RASSF4* (Ras Association Domain Family Member 4), *SPG21* (SPG21 Abhydrolase Domain Containing, Maspardin, involved in repression of T cell activation (Soderblom et al., 2010)), *MYOG* (Myogenin) were significantly differentially expressed in young myoblasts, *TCF4* (Transcription Factor 4), *MTPN* (Myotrophin), *SAMD1* (Sterile Alpha Motif Domain Containing 1, an unmethylated CGI-binding protein (Stielow et al., 2021)) and *MAB21L1* (Mab-21 Like 1, a putative nucleoidyltransferase (Rad et al., 2019)) in adult myoblasts and *MYBPC1* (Myosin Binding protein C1), *TNNC2* (Troponin C2, fast skeletal type), *TPM1* (Tropomysin) and *TBX3* (T-Box Transcription Factor 3) in aged myoblasts. Mitochondrial translation terms were enriched in young myoblasts while muscle processes, differentiation and development were enriched in aged myoblast (**Figure 4B** and **figure 4-figure supplement 2**).

**Figure 4:**
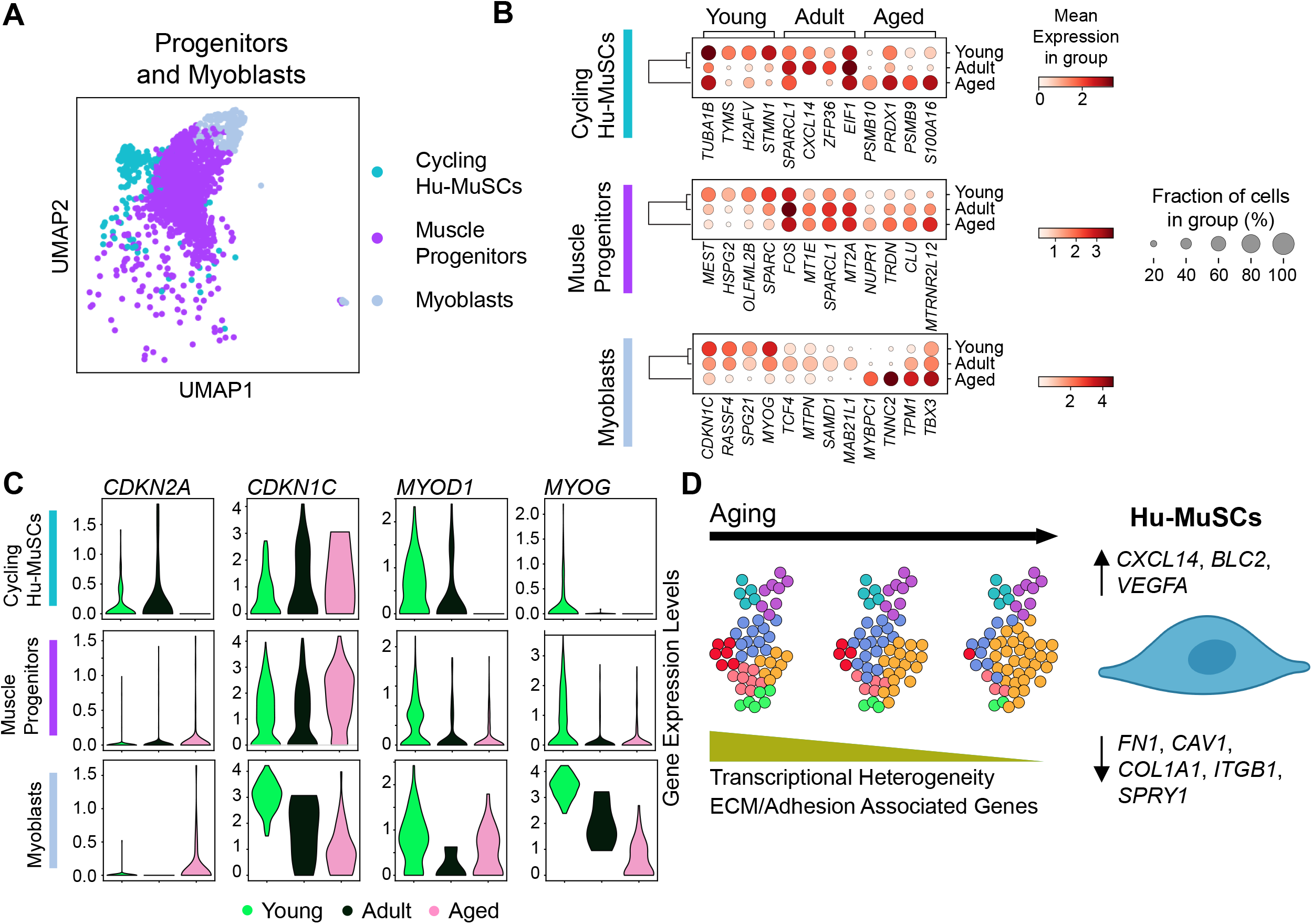
Transcriptional analysis of progenitor and myoblast cells in the different age groups. (A) UMAP of merged age groups for only muscle progenitors (cluster 4), cycling Hu-MuSCs (cluster 8) and Myoblasts (cluster 9). (B) Dot plot displaying the top 4 genes differentially expressed for each age group in cluster 4, 8 and 9. (C) Violin plots of the expression of genes altered with aging. (D) Summary schematic of how Hu-MuSCs transcriptome changes with aging.

In addition, we found that *CDKN2A* (Cyclin Dependent Kinase Inhibitor 2A) and *CDKN1C* expression increased with aging in cycling Hu-MuSCs and muscle progenitor cells. In contrast, MYOD1 and *MYOG* levels decreased with aging in those two clusters as well as in the myoblast cluster. *CDKN2A* transcription levels also increased in myoblasts upon aging while *CDKN1C* levels decreased in the myoblast cluster (**Figure 4C**).

Overall, these analyses provide new insights into transcriptomic modulations of more differentiated human muscle stem cells and muscle progenitors during aging summarily characterized by similar age-related GO term enrichment to Hu-MuSCs.

## DISCUSSION

In this extension of our prior work we performed extended transcriptomic analyses of isolated human muscle stem cells across a range of adult ages. We found that satellite cell aging is characterized by a global decrease in transcriptional heterogeneity. At the single gene level, new and previously described transcriptional age-related changes in human satellite cells were identified.

The addition of new samples coupled with more complex computational analyses of our single cell RNA sequencing data allowed us to further understand the cellular heterogeneity of human satellite cells. While aging was not associated with the appearance or disappearance of age-specific clusters, we found that the cell distribution among the different clusters was altered. A loss of cellular heterogeneity during mouse muscle aging has been shown previously, Chakkalakal et al. found a decrease of labeling retaining-SCs (bone fide stem cells) while committed progenitors (non-labeling retaining-SCs) were preserved in aged mice (Chakkalakal et al., 2012). This suggests that certain subpopulations of Hu-MuSCs are retained with aging while others are reduced or partially lost. Importantly, this loss of heterogeneity was associated with a decline in transplantation potential of aged SCs (Chakkalakal et al., 2012). These prior studies together with this study suggest that a shift in satellite cell subpopulation representation may be responsible for impaired muscle regeneration in the aging population.

We found (1) the distribution of cells among the various human clusters we have characterized changes during aging and (2) genes including *CAV1, CXCL14, FN1* and *GPX3* that can explain this differential cell distribution. Our analysis highlighted gene-defining clusters that are significantly altered by aging. For example, we found that *CAV1*, a marker of a high transplantation potential Hu-MuSC subpopulation (Barruet et al., 2020), decreases with age. We (Barruet et al., 2020) and others (Baker & Tuan, 2013) have previously described that *CAV1* is expressed in quiescent satellite cells with increased engraftment properties, although its functional role in aging satellite cells remains unknown. Indeed, CAV1 in aging is not well understood as opposing finding a have been described in several tissues and studies (Gurley, Standifer, & Hargis, 2021; Ha et al., 2021; Head et al., 2010; Kruglikov, Zhang, & Scherer, 2019; Wicher, Prakash, & Pabelick, 2019). Further functional studies in human SCs are necessary to determine the precise role of *CAV1* in aging and future transplantation studies will help determine if diminishment of this satellite cell subpopulation is responsible for decreased satellite cell function in aging.

CXCL14, transcripts of which we found increased in aging, has been shown to prevent cell cycle withdrawal and to be a negative regulator of myoblast differentiation. Experimental CXCL14 reduction ameliorates regenerative defects in aging mouse muscle (Waldemer-Streyer et al., 2017). We also found an increase of *GPX3* expression in aged Hu-MuSCs. GPX3 (glutathione peroxidase 3), a retinoid-responsive gene that mediates the antioxidant effects of retinoic acid in human myoblasts, may be important in muscle stem cell survival (El Haddad et al., 2012). Therefore, our observed increase in *CXCL14+* and *GPX3+* Hu-MuSCs could be related to regenerative decline of aged human muscles.

Our in-depth computational analyses of human samples also corroborated other trends previously described in age-related mouse studies, notably a decrease in *ITGB1* and *SPRY1* expression in aging. Levels of ITGB1 and FAK (Focal Adhesion Kinase) were lower in aged satellite cells leading to decreased cell adhesion and increased cell death (Lukjanenko et al., 2016; Rozo et al., 2016). In addition to a decrease of ITGB1 and FAK, our analyses also reveal a decrease in ECM such as fibronectin and collagen associated genes. While extensive work has been done on ECM remodeling of the satellite cell’s niche during aging (Evano & Tajbakhsh, 2018), there is a lack of data describing the effect of aging on ECM components produced by satellite cells. Nevertheless, fibronectin and collagens produced by satellite cells are critical for the maintenance of their quiescence (Baghdadi et al., 2018; Bentzinger et al., 2013). Our analyses suggest that aging may induce a decrease in ECM components expression by Hu-MuSCs which may have a role in ECM remodeling found in aging muscles (Schüler et al., 2021).

SPRY1, a regulator of satellite cell return to quiescence (Shea et al., 2010) and detected in a subset of Hu-MuSCs in our previous (Barruet et al., 2020) and present studies, also decreases in aged SCs. Comparable results were found in mice, where age-associated methylation suppression of SPRY1 leads to loss of the reserve stem cell pool (Bigot et al., 2015) while SPRY1 over-expression in aged satellite cells in vivo preserves the SC pool (Chakkalakal et al., 2012). Our findings add additional evidence supporting the concept that the high expressing *SPRY1* subset of SCs is critical for muscle regeneration during aging. We also found increased expression of other major drivers of ageing such as mTor pathway targets (e.i. *BCL2* and *VEGFA)* (Liu & Sabatini, 2020) being elevated in aged satellite cells. FOS was also found to be elevated in cluster 1 where most aged cells resided. Although, a recent study showed that Fos mRNA is a feature of freshly isolated satellite cells from uninjured muscle and that it marks a subset of satellite cells with enhanced regenerative ability (Almada et al., 2021), how FOS levels impact aging in human muscle stem cells still remains to be fully elucidated. Finally, our dataset also captured more differentiated muscle stem cells in which *CDKN2A* expression level was increased and *MYOD1* and *MYOG* levels were decreased with age. Indeed, increased level of *CDKN2A* has been described in geriatric human and mouse muscle stem cells to induce a loss of reversible quiescence, a pre-senescence state and result in failure to proliferate and differentiate (Sousa-Victor et al., 2014). Altogether, with a limited number of samples, our human satellite cell transcriptomic study was able to validate age-related mouse findings, confirm the potential role of age-related pathways in Hu-MuSCs during aging, and identify changes in satellite cell distribution among subpopulations.

Our RNA velocity analysis identified additional genes with age-specific differential velocity which explained the vector field of the different Hu-MuSCs age groups. While some of those will need further experimental investigation to understand their mechanism of action during aging (e.g *CDKN1C, DIO2, KLF6, NPTX2, MAP1B*) other genes, for which rodent experiments, have been carried out in homeostasis, may explain the loss of satellite cell stemness and self-renewal such as *GREM1* and *EDN3. GREM1* velocity was associated with aged Hu-MuSCs. A BMP antagonist, GREM1 would be expected to induce a decrease in satellite cell number (Borok et al., 2020) possibly by acting as a negative regulator of satellite cell self-renewal. Therefore GREM1 could account for the loss of self-renewal and reduced number of satellite cells in aged human muscle. Separately, we found that *EDN3* explains the vector field of young Hu-MuSCs. *EDN3* is expressed in quiescent mouse satellite cells (Fukada et al., 2007) and is down-regulated as they become activated (Machado et al., 2017; van Velthoven, de Morree, Egner, Brett, & Rando, 2017) suggesting that EDN3 could play a functional role and/or be a new marker for the loss of quiescence with aging.

This study is the first report of single cell transcriptomes of human satellite cells at various stages of aging. The possibility exists that our representation of aging human satellite cell transcriptomes is incomplete with a limited number of samples. However, since we were able to confirm previous mouse observations, it is likely that this study does contain a faithful and adequate sampling of human muscles to describe the major alterations of the transcriptomic landscape in aging. In situ validations and functional studies will further elucidate the roles of the different genes identified here.

## MATERIALS AND METHODS

### Key Resources Table

| Key Resources Table |  |  |  |  |
| --- | --- | --- | --- | --- |
| Reagent type (species) or resource | Designation | Source or reference | Identifiers | Additional information |
| software, algorithm | Kallisto/loompy 3.0 package | https://linnarssonlab.org/loompy/kallisto/index.html | SCR_016666 |  |
| software, algorithm | Scanpy version 1.9 | https://scanpy.readthedocs.io/en/latest/tutorials.html | SCR_018139 |  |
| software, algorithm | scVelo v0.2.4 | https://scvelo.readthedocs.io | SCR_018168 |  |

### Human Specimen Procurement and Hu-MuSCs isolation

This study was conducted under the approval of the Institutional Review Board at The University of California San Francisco (UCSF). Biopsies were obtained from individuals undergoing surgery at UCSF. Written informed consent was obtained from all subjects. All types of muscle used for each experiment are listed in **Figure 1-Source Data 1**. Additional CXCR4+/CD29+/CD65+ Hu-MuSC samples were isolated as described in (Barruet et al., 2020; Garcia et al., 2018; Garcia, Tamaki, Xu, & Pomerantz, 2017; X. Xu et al., 2015).

### Single cell RNA Sequencing and Analysis

Single cell RNA sequencing and gene core matrices retrieval of additional samples were performed as described in (Barruet et al., 2020). Gene-barcoded matrices were analyzed with the Python package Scanpy version 1.9 (F. Alexander Wolf et al., 2018). For each sample, cells with fewer than 500 genes, greater than 7000 genes and genes expressed in fewer than 5 cells were not included in the downstream analyses. Cells with more than 15% mitochondrial counts were filtered out. Each sample was first merged into its own age group with batch balanced k nearest neighbor (BBKNN) algorithm (Polański et al., 2019) to remove potential technical variation between samples. A resolution of 0.5 was used for all subset age group. Cluster were annotated using known markers found in the literature combined with differentially expressed genes (Wilcoxon test, function sc.tl.rank_genes_groups). Since after filtering the adult group contained 66,905 cells, and the young and aged group contained 11,502 cells and 9,407 cells respectively, to avoid cofounding factors due to discrepancy in cell number among the three age groups, we downsampled the adult cell group to 11,000 cells using the sc.pp.subsample function. Following this, the ‘adult’ and ‘aged’ datasets were integrated onto the annotated ‘young’ dataset using the Scanpy INGEST function sc.tl.ingest. Differential expression analysis was performed between the age groups using the same Wilcoxon statistical test, as implemented in Scanpy. Marker gene expression was visualized using either dot-plots, where the size of the dot reflects the percentage of cells expressing the gene and the color indicated the relative expression, or violin plots, with the width of the violin plot depicting the larger probability density of cells expressing each gene at the indicated expression levels.

### scVelo and PAGA trajectory analysis

Count matrices (unspliced) and mature (spliced) abundances were generated for each sample from fastq files using Kallisto/loompy 3.0 package. scVelo v0.2.4 package implemented into Scanpy was used to perform RNA velocity analysis (Bergen et al., 2020). Datasets were processed using the recommended parameters as described in Scanpy scVelo implementation (Bergen et al., 2020). The age group samples were pre-processed using scv.pp.filter and scv.pp.normalize followed by scv.pp.moments functions for detection of minimum number of counts, filtering and normalization. scv.tl.velocity and scv.tl.velocity_graph functions were used to calculate and visualized gene specific velocities. Gene ranking for each age group resulting from differential velocity t-test was perform using the scv.tl.rank_velocity_genes. Scanpy implemented partition-based graph abstraction (PAGA) functions (scv.tl.paga and scv.pl.paga) was used to assess the data topology with weighted edges corresponding to the connectivity between two clusters. Default parameters were used (F. A. Wolf et al., 2019).

### Gene Ontology analysis

Differentially expressed genes, p-values and fold changes were used as input to generate GO-term enrichment with the clusterProfiler package in R. Thresholds were set p-value <0.05 and fold change >1 for the GO-term analysis.

## Data and materials availability

Single cell gene expression fastq files and filtered matrices have been deposited (GSE196554). Detailed scripts can be found here, https://github.com/EmilieB12/Aging_Hu-MuSCs.

## ACKNOWLEDGEMENTS

This work was supported by NIH R01AR072638-03 to JHP. The authors would like to thank all the organ and tissue donors and their families for their generous donation.

---

**Figure 1-figure supplement 1:**
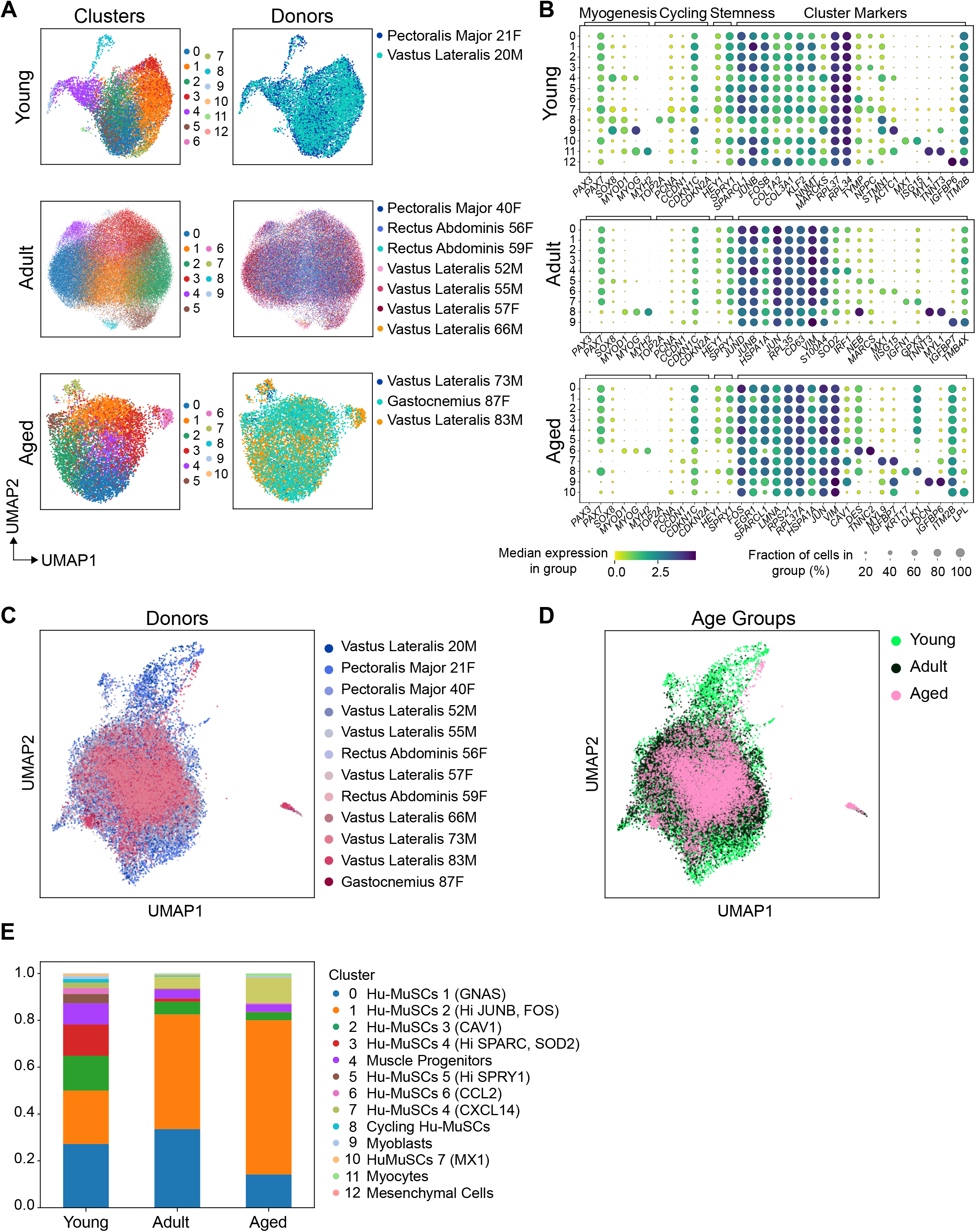
BBKNN and INGEST merging of sorted human muscle stem cells. (A) As shown in Figure 1A, samples were first merged into their own age group. UMAP displaying clusters and samples for each age group. (B) Dot plots displaying the expression of myogenic, cycling, stemness and cluster marker genes for each cluster in each age group. (C) UMAP of the INGEST analysis displaying all 12 samples. (D) MAP of each age group and their distribution in clusters. (E) Proportion plot of cells assigned to each cluster for each age group.

**Figure 2-figure supplement 1:**
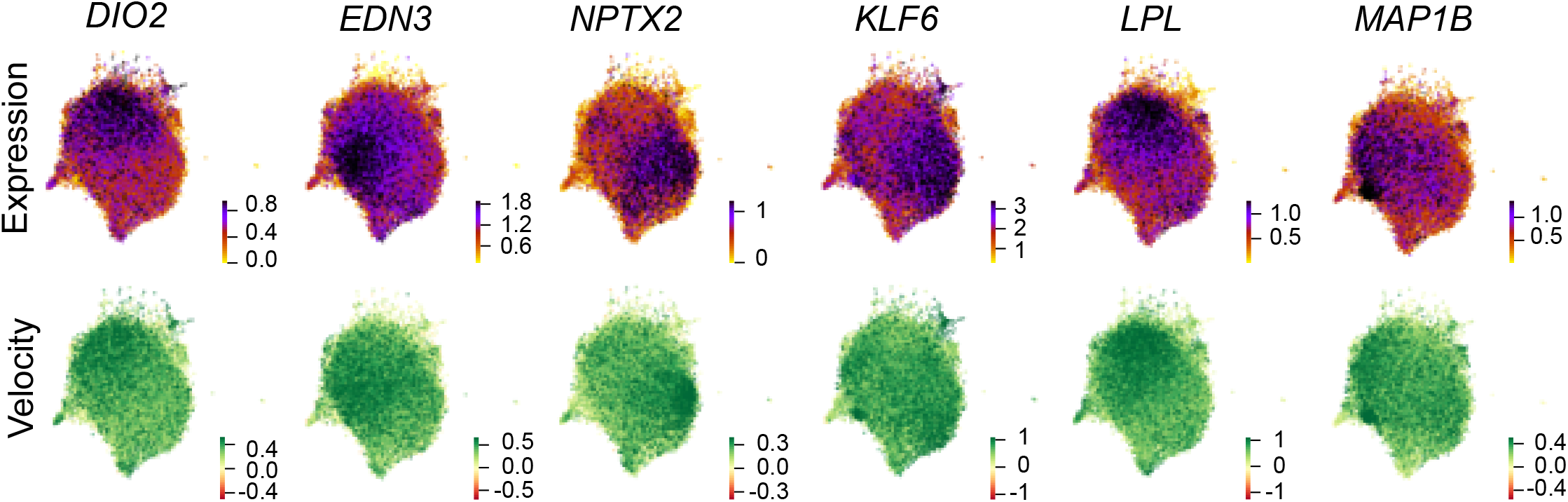
Expression and velocity of relevant genes that inferred age-specific differential velocity.

**Figure 4-figure supplement 1:**
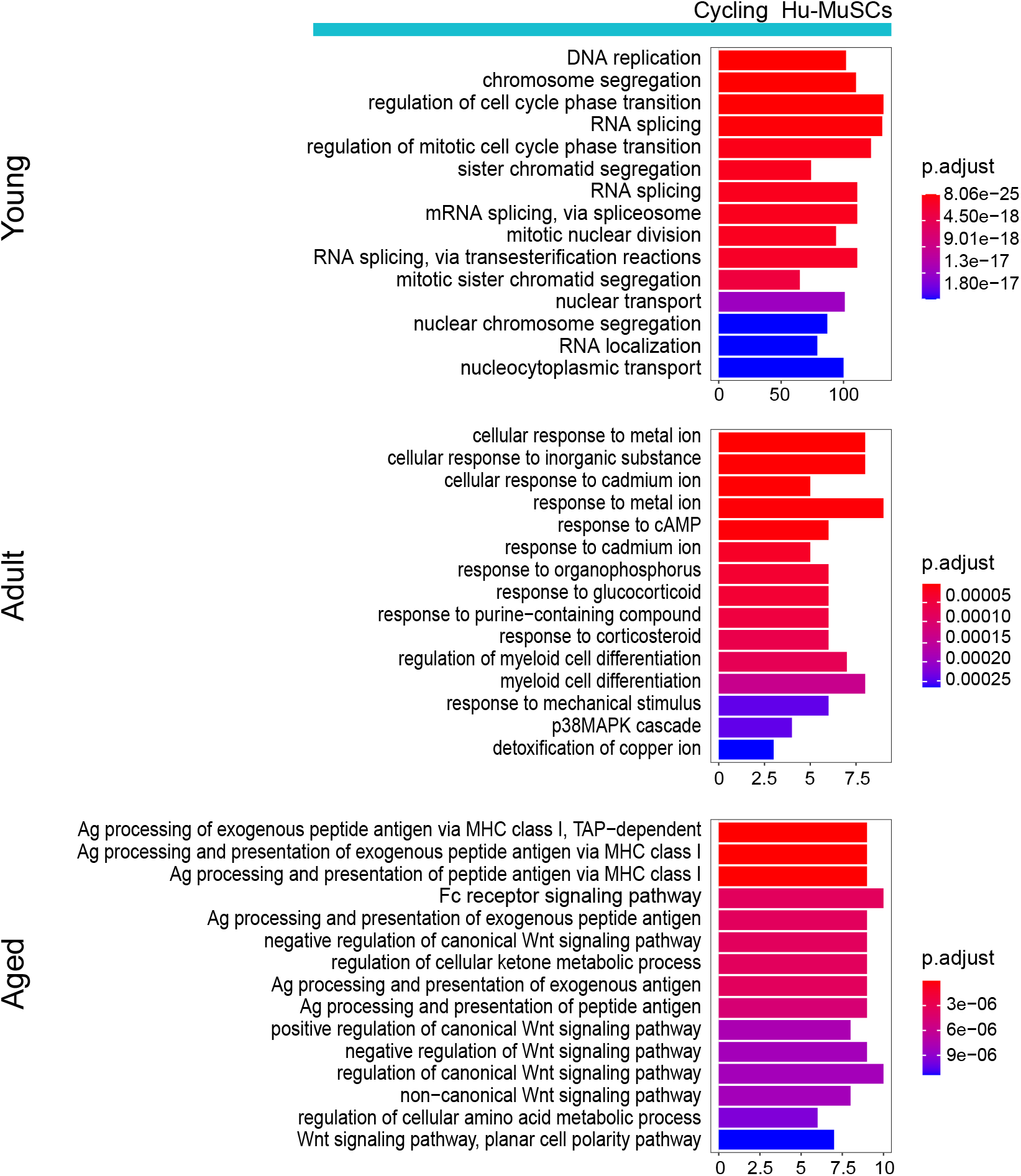
Cycling Hu-MuSCs GO term analysis upon aging. Bar plots of gene ontology analysis of differentially up-regulated genes in the cycling Hu-MuSCs cluster (8) for each age group.

**Figure 4-figure supplement 2:**
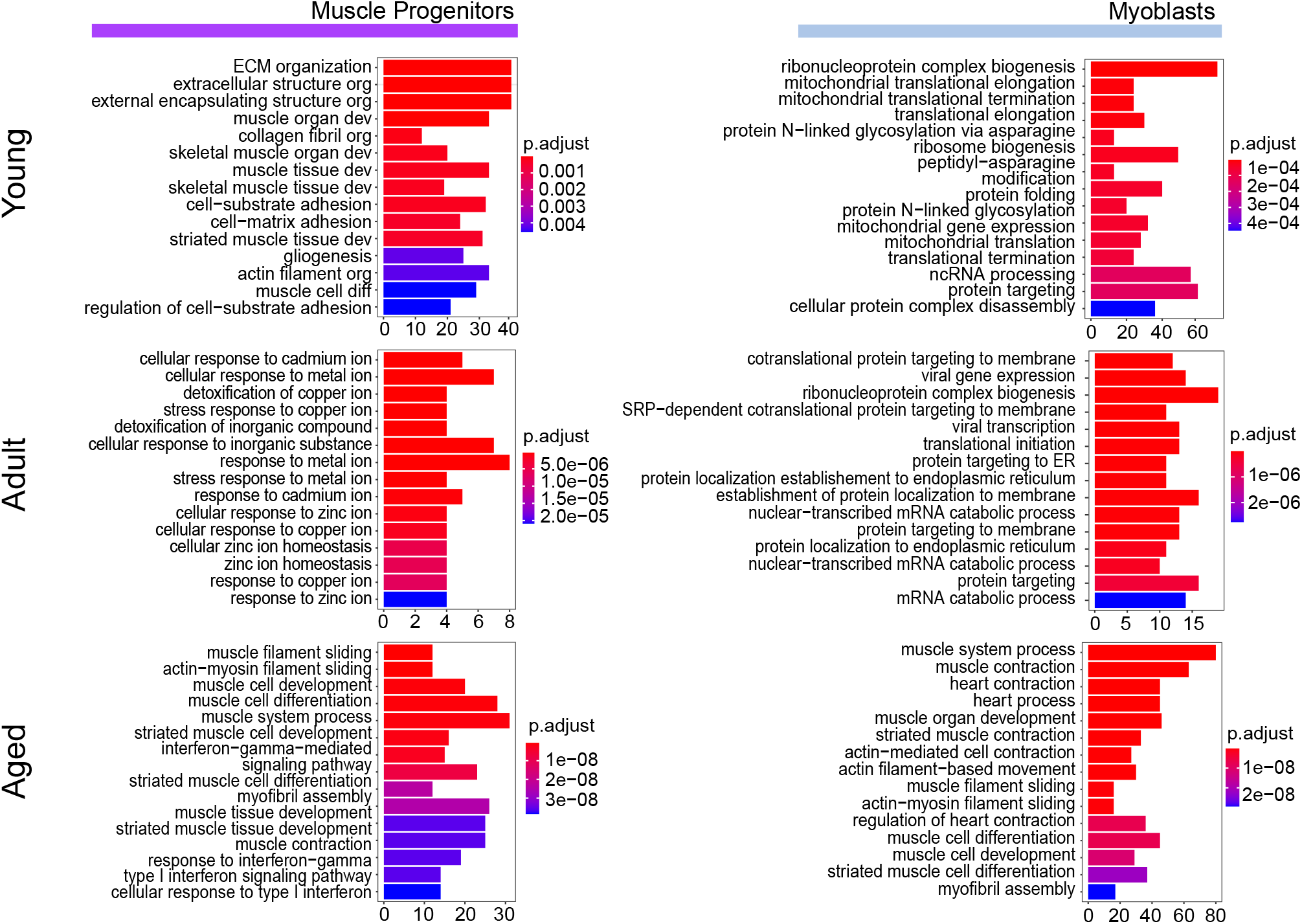
Progenitor and myoblast cells GO term analysis upon aging. Bar plots of gene ontology analysis of differentially up-regulated genes in the muscle progenitor cluster (4), cycling Hu-MuSCs cluster (8) and Myoblasts cluster (9) for each age group.

## ADDITIONAL FILES

**Figure 1-Source Data 1: Demographics and sample characteristics collected and used for downstream analysis.**

**Figure 2-Source Data 1: Gene ranking for each age group resulting from differential velocity t-test.**

## Notes

### Competing Interest Statement

The authors have declared no competing interest.

## REFERENCES

Almada, A. E., Horwitz, N., Price, F. D., Gonzalez, A. E., Ko, M., Bolukbasi, O. V.,… Wagers, A. J. (2021). FOS licenses early events in stem cell activation driving skeletal muscle regeneration. Cell Rep, 34(4), 108656. doi:10.1016/j.celrep.2020.108656

Angelidis, I., Simon, L. M., Fernandez, I. E., Strunz, M., Mayr, C. H., Greiffo, F. R.,… Strom, T.-M. (2019). An atlas of the aging lung mapped by single cell transcriptomics and deep tissue proteomics. Nature communications, 10(1), 1–17.

Arpke, R. W., Shams, A. S., Collins, B. C., Larson, A. A., Lu, N., Lowe, D. A., & Kyba, M. (2021). Preservation of satellite cell number and regenerative potential with age reveals locomotory muscle bias. Skeletal Muscle, 11(1), 22. doi:10.1186/s13395-021-00277-2

Baghdadi, M. B., Castel, D., Machado, L., Fukada, S.-i., Birk, D. E., Relaix, F.,… Mourikis, P. (2018). Reciprocal signalling by Notch–Collagen V–CALCR retains muscle stem cells in their niche. Nature, 557(7707), 714–718. doi:10.1038/s41586-018-0144-9

Baker, N., & Tuan, R. S. (2013). The less-often-traveled surface of stem cells: caveolin-1 and caveolae in stem cells, tissue repair and regeneration. Stem Cell Res Ther, 4(4), 90. doi:10.1186/scrt276

Barruet, E., Garcia, S. M., Striedinger, K., Wu, J., Lee, S., Byrnes, L.,… Pomerantz, J. H. (2020). Functionally heterogeneous human satellite cells identified by single cell RNA sequencing. Elife, 9. doi:10.7554/eLife.51576

Bentzinger, C. F., Wang, Y. X., von Maltzahn, J., Soleimani, V. D., Yin, H., & Rudnicki, M. A. (2013). Fibronectin regulates Wnt7a signaling and satellite cell expansion. Cell Stem Cell, 12(1), 75–87. doi:10.1016/j.stem.2012.09.015

Bergen, V., Lange, M., Peidli, S., Wolf, F. A., & Theis, F. J. (2020). Generalizing RNA velocity to transient cell states through dynamical modeling. Nature Biotechnology, 38(12), 1408–1414. doi:10.1038/s41587-020-0591-3

Bigot, A., Duddy, W. J., Ouandaogo, Z. G., Negroni, E., Mariot, V., Ghimbovschi, S.,… Duguez, S. (2015). Age-Associated Methylation Suppresses SPRY1, Leading to a Failure of Re-quiescence and Loss of the Reserve Stem Cell Pool in Elderly Muscle. Cell Reports, 13(6), 1172–1182. doi:https://doi.org/10.1016/j.celrep.2015.09.067

Bik-Multanowski, M., Pietrzyk, J. J., & Midro, A. (2015). MTRNR2L12: A Candidate Blood Marker of Early Alzheimer’s Disease-Like Dementia in Adults with Down Syndrome. J Alzheimers Dis, 46(1), 145–150. doi:10.3233/jad-143030

Borok, M. J., Mademtzoglou, D., & Relaix, F. (2020). Bu-M-P-ing Iron: How BMP Signaling Regulates Muscle Growth and Regeneration. J Dev Biol, 8(1). doi:10.3390/jdb8010004

Brack, A. S., & Muñoz-Cánoves, P. (2016). The ins and outs of muscle stem cell aging. Skeletal Muscle, 6(1), 1. doi:10.1186/s13395-016-0072-z

Buroker, N. E. (2014). Regulatory SNPs and transcriptional factor binding sites in ADRBK1, AKT3, ATF3, DIO2, TBXA2R and VEGFA. Transcription, 5(4), e964559. doi:10.4161/21541264.2014.964559

Chakkalakal, J. V., Jones, K. M., Basson, M. A., & Brack, A. S. (2012). The aged niche disrupts muscle stem cell quiescence. Nature, 490(7420), 355–360. doi:10.1038/nature11438

Chang, N. C., & Rudnicki, M. A. (2014). Satellite cells: the architects of skeletal muscle. Curr Top Dev Biol, 107, 161–181. doi:10.1016/b978-0-12-416022-4.00006-8

Charville, G. W., Cheung, T. H., Yoo, B., Santos, P. J., Lee, G. K., Shrager, J. B., & Rando, T. A. (2015). Ex Vivo Expansion and In Vivo Self-Renewal of Human Muscle Stem Cells. Stem Cell Reports, 5(4), 621–632. doi:10.1016/j.stemcr.2015.08.004

Conboy, I. M., Conboy, M. J., Smythe, G. M., & Rando, T. A. (2003). Notch-mediated restoration of regenerative potential to aged muscle. Science, 302(5650), 1575–1577. doi:10.1126/science.1087573

Consortium, T. M. (2020). A single cell transcriptomic atlas characterizes aging tissues in the mouse. Nature, 583(7817), 590.

Curtis, E., Litwic, A., Cooper, C., & Dennison, E. (2015). Determinants of Muscle and Bone Aging. J Cell Physiol, 230(11), 2618–2625. doi:10.1002/jcp.25001

De Santis, R., Etoc, F., Rosado-Olivieri, E. A., & Brivanlou, A. H. (2021). Self-organization of human dorsal-ventral forebrain structures by light induced SHH. Nature communications, 12(1), 6768. doi:10.1038/s41467-021-26881-w

Ding, C., Fan, X., & Wu, G. (2017). Peroxiredoxin 1 - an antioxidant enzyme in cancer. J Cell Mol Med, 21(1), 193–202. doi:10.1111/jcmm.12955

Distefano, G., & Goodpaster, B. H. (2018). Effects of Exercise and Aging on Skeletal Muscle. Cold Spring Harb Perspect Med, 8(3). doi:10.1101/cshperspect.a029785

El Haddad, M., Jean, E., Turki, A., Hugon, G., Vernus, B., Bonnieu, A.,… Carnac, G. (2012). Glutathione peroxidase 3, a new retinoid target gene, is crucial for human skeletal muscle precursor cell survival. Journal of Cell Science, 125(24), 6147–6156. doi:10.1242/jcs.115220

Ermolaeva, M., Neri, F., Ori, A., & Rudolph, K. L. (2018). Cellular and epigenetic drivers of stem cell ageing. Nature Reviews Molecular Cell Biology, 19(9), 594–610. doi:10.1038/s41580-018-0020-3

Evano, B., & Tajbakhsh, S. (2018). Skeletal muscle stem cells in comfort and stress. npj Regenerative Medicine, 3(1), 24. doi:10.1038/s41536-018-0062-3

Fukada, S., Uezumi, A., Ikemoto, M., Masuda, S., Segawa, M., Tanimura, N.,… Takeda, S. (2007). Molecular signature of quiescent satellite cells in adult skeletal muscle. Stem Cells, 25(10), 2448–2459. doi:10.1634/stemcells.2007-0019

Garcia, S. M., Tamaki, S., Lee, S., Wong, A., Jose, A., Dreux, J.,… Pomerantz, J. H. (2018). High-Yield Purification, Preservation, and Serial Transplantation of Human Satellite Cells. Stem Cell Reports, 10(3), 1160–1174. doi:10.1016/j.stemcr.2018.01.022

Garcia, S. M., Tamaki, S., Xu, X., & Pomerantz, J. H. (2017). Human Satellite Cell Isolation and Xenotransplantation. Methods Mol Biol, 1668, 105–123. doi:10.1007/978-1-4939-7283-8_8

García-Prat, L., Martínez-Vicente, M., Perdiguero, E., Ortet, L., Rodríguez-Ubreva, J., Rebollo, E.,… Serrano, A. L. (2016). Autophagy maintains stemness by preventing senescence. Nature, 529(7584), 37–42.

Gurley, J., Standifer, N., & Hargis, E. A. (2021). Progressive loss of retinal arteriolar smooth muscle cells (SMCs) with aging and caveolin-1 (Cav1) depletion: Assessing the importance of endothelial cell (EC)-Cav1 in retinal SMC maintenance. Investigative Ophthalmology & Visual Science, 62(8), 2728–2728.

Ha, T.-Y., Choi, Y. R., Noh, H. R., Cha, S.-H., Kim, J.-B., & Park, S. M. (2021). Age-related increase in caveolin-1 expression facilitates cell-to-cell transmission of α-synuclein in neurons. Molecular Brain, 14(1), 122. doi:10.1186/s13041-021-00834-2

Halpain, S., & Dehmelt, L. (2006). The MAP1 family of microtubule-associated proteins. Genome Biol, 7(6), 224. doi:10.1186/gb-2006-7-6-224

Head, B. P., Peart, J. N., Panneerselvam, M., Yokoyama, T., Pearn, M. L., Niesman, I. R.,… Patel, H. H. (2010). Loss of Caveolin-1 Accelerates Neurodegeneration and Aging. PLOS ONE, 5(12), e15697. doi:10.1371/journal.pone.0015697

Jørgensen, L. H., Jepsen, P. L., Boysen, A., Dalgaard, L. B., Hvid, L. G., Ørtenblad, N.,… Schrøder, H. D. (2017). SPARC Interacts with Actin in Skeletal Muscle in Vitro and in Vivo. Am J Pathol, 187(2), 457–474. doi:10.1016/j.ajpath.2016.10.013

Karbiener, M., Glantschnig, C., Pisani, D. F., Laurencikiene, J., Dahlman, I., Herzig, S.,… Scheideler, M. (2015). Mesoderm-specific transcript (MEST) is a negative regulator of human adipocyte differentiation. International Journal of Obesity, 39(12), 1733–1741. doi:10.1038/ijo.2015.121

Kawanabe, Y., & Nauli, S. M. (2011). Endothelin. Cell Mol Life Sci, 68(2), 195–203. doi:10.1007/s00018-010-0518-0

Kimmel, J. C., Penland, L., Rubinstein, N. D., Hendrickson, D. G., Kelley, D. R., & Rosenthal, A. Z. (2019). Murine single-cell RNA-seq reveals cell-identity-and tissue-specific trajectories of aging. Genome research, 29(12), 2088–2103.

Kimmel, J. C., Yi, N., Roy, M., Hendrickson, D. G., & Kelley, D. R. (2021). Differentiation reveals latent features of aging and an energy barrier in murine myogenesis. Cell Reports, 35(4), 109046. doi:https://doi.org/10.1016/j.celrep.2021.109046

Kowalczyk, M. S., Tirosh, I., Heckl, D., Rao, T. N., Dixit, A., Haas, B. J.,… Regev, A. (2015). Single-cell RNA-seq reveals changes in cell cycle and differentiation programs upon aging of hematopoietic stem cells. Genome research, 25(12), 1860–1872.

Kruglikov, I. L., Zhang, Z., & Scherer, P. E. (2019). Caveolin-1 in skin aging – From innocent bystander to major contributor. Ageing Research Reviews, 55, 100959. doi:https://doi.org/10.1016/j.arr.2019.100959

La Manno, G., Soldatov, R., Zeisel, A., Braun, E., Hochgerner, H., Petukhov, V.,… Kharchenko, P. V. (2018). RNA velocity of single cells. Nature, 560(7719), 494–498. doi:10.1038/s41586-018-0414-6

Lee, D., Takayama, S., & Goldberg, A. L. (2018). ZFAND5/ZNF216 is an activator of the 26S proteasome that stimulates overall protein degradation. Proc Natl Acad Sci U S A, 115(41), E9550–e9559. doi:10.1073/pnas.1809934115

Liu, G. Y., & Sabatini, D. M. (2020). mTOR at the nexus of nutrition, growth, ageing and disease. Nature Reviews Molecular Cell Biology, 21(4), 183–203. doi:10.1038/s41580-019-0199-y

Liu, J., Li, J., Wang, K., Liu, H., Sun, J., Zhao, X.,… Yang, A. (2021). Aberrantly high activation of a FoxM1–STMN1 axis contributes to progression and tumorigenesis in FoxM1-driven cancers. Signal Transduction and Targeted Therapy, 6(1), 42. doi:10.1038/s41392-020-00396-0

Liu, J., Song, X., Kuang, F., Zhang, Q., Xie, Y., Kang, R.,… Tang, D. (2021). NUPR1 is a critical repressor of ferroptosis. Nat Commun, 12(1), 647. doi:10.1038/s41467-021-20904-2

Lukjanenko, L., Jung, M. J., Hegde, N., Perruisseau-Carrier, C., Migliavacca, E., Rozo, M.,… Bentzinger, C. F. (2016). Loss of fibronectin from the aged stem cell niche affects the regenerative capacity of skeletal muscle in mice. Nature Medicine, 22(8), 897–905. doi:10.1038/nm.4126

Machado, L., Esteves de Lima, J., Fabre, O., Proux, C., Legendre, R., Szegedi, A.,… Mourikis, P. (2017). In Situ Fixation Redefines Quiescence and Early Activation of Skeletal Muscle Stem Cells. Cell Rep, 21(7), 1982–1993. doi:10.1016/j.celrep.2017.10.080

Mademtzoglou, D., Asakura, Y., Borok, M. J., Alonso-Martin, S., Mourikis, P., Kodaka, Y.,… Relaix, F. (2018). Cellular localization of the cell cycle inhibitor Cdkn1c controls growth arrest of adult skeletal muscle stem cells. Elife, 7. doi:10.7554/eLife.33337

Martinez, J. R., Dhawan, A., & Farach-Carson, M. C. (2018). Modular Proteoglycan Perlecan/HSPG2: Mutations, Phenotypes, and Functions. Genes (Basel), 9(11). doi:10.3390/genes9110556

Marty, I., & Fauré, J. (2016). Excitation-Contraction Coupling Alterations in Myopathies. J Neuromuscul Dis, 3(4), 443–453. doi:10.3233/jnd-160172

Polański, K., Young, M. D., Miao, Z., Meyer, K. B., Teichmann, S. A., & Park, J.-E. (2019). BBKNN: fast batch alignment of single cell transcriptomes. Bioinformatics, 36(3), 964–965. doi:10.1093/bioinformatics/btz625

Rad, A., Altunoglu, U., Miller, R., Maroofian, R., James, K. N., Çağlayan, A. O.,… Schmidts, M. (2019). MAB21L1 loss of function causes a syndromic neurodevelopmental disorder with distinctive <em>c</em>erebellar, <em>o</em>cular, cranio<em>f</em>acial and <em>g</em>enital features (COFG syndrome). Journal of Medical Genetics, 56(5), 332–339. doi:10.1136/jmedgenet-2018-105623

Rozo, M., Li, L., & Fan, C.-M. (2016). Targeting β1-integrin signaling enhances regeneration in aged and dystrophic muscle in mice. Nature Medicine, 22(8), 889–896. doi:10.1038/nm.4116

Scaramozza, A., Park, D., Kollu, S., Beerman, I., Sun, X., Rossi, D. J.,… Brack, A. S. (2019). Lineage Tracing Reveals a Subset of Reserve Muscle Stem Cells Capable of Clonal Expansion under Stress. Cell Stem Cell, 24(6), 944–957 e945. doi:10.1016/j.stem.2019.03.020

Schäfer, R., Zweyer, M., Knauf, U., Mundegar, R. R., & Wernig, A. (2005). The ontogeny of soleus muscles in mdx and wild type mice. Neuromuscul Disord, 15(1), 57–64. doi:10.1016/j.nmd.2004.09.011

Schüler, S. C., Kirkpatrick, J. M., Schmidt, M., Santinha, D., Koch, P., Di Sanzo, S.,… von Maltzahn, J. (2021). Extensive remodeling of the extracellular matrix during aging contributes to age-dependent impairments of muscle stem cell functionality. Cell Rep, 35(10), 109223. doi:10.1016/j.celrep.2021.109223

Shea, K. L., Xiang, W., LaPorta, V. S., Licht, J. D., Keller, C., Basson, M. A., & Brack, A. S. (2010). Sprouty1 regulates reversible quiescence of a self-renewing adult muscle stem cell pool during regeneration. Cell Stem Cell, 6(2), 117–129. doi:10.1016/j.stem.2009.12.015

Shi, N., Guo, X., & Chen, S. Y. (2014). Olfactomedin 2, a novel regulator for transforming growth factor-β-induced smooth muscle differentiation of human embryonic stem cell-derived mesenchymal cells. Mol Biol Cell, 25(25), 4106–4114. doi:10.1091/mbc.E14-08-1255

Snijders, T., Nederveen, J. P., McKay, B. R., Joanisse, S., Verdijk, L. B., van Loon, L. J. C., & Parise, G. (2015). Satellite cells in human skeletal muscle plasticity. Frontiers in Physiology, 6. doi:10.3389/fphys.2015.00283

Soderblom, C., Stadler, J., Jupille, H., Blackstone, C., Shupliakov, O., & Hanna, M. C. (2010). Targeted disruption of the Mast syndrome gene SPG21 in mice impairs hind limb function and alters axon branching in cultured cortical neurons. Neurogenetics, 11(4), 369–378. doi:10.1007/s10048-010-0252-7

Sousa-Victor, P., Gutarra, S., García-Prat, L., Rodriguez-Ubreva, J., Ortet, L., Ruiz-Bonilla, V.,… Muñoz-Cánoves, P. (2014). Geriatric muscle stem cells switch reversible quiescence into senescence. Nature, 506(7488), 316–321. doi:10.1038/nature13013

Stielow, B., Zhou, Y., Cao, Y., Simon, C., Pogoda, H.-M., Jiang, J.,… Liefke, R. (2021). The SAM domain-containing protein 1 (SAMD1) acts as a repressive chromatin regulator at unmethylated CpG islands. Science Advances, 7(20), eabf2229. doi:doi:10.1126/sciadv.abf2229

Stuart, T., Butler, A., Hoffman, P., Hafemeister, C., Papalexi, E., Mauck, W. M., 3rd,… Satija, R. (2019). Comprehensive Integration of Single-Cell Data. Cell, 177(7), 1888–1902.e1821. doi:10.1016/j.cell.2019.05.031

Tetreault, M. P., Yang, Y., & Katz, J. P. (2013). Krüppel-like factors in cancer. Nat Rev Cancer, 13(10), 701–713. doi:10.1038/nrc3582

van Velthoven, C. T. J., de Morree, A., Egner, I. M., Brett, J. O., & Rando, T. A. (2017). Transcriptional Profiling of Quiescent Muscle Stem Cells In Vivo. Cell Rep, 21(7), 1994–2004. doi:10.1016/j.celrep.2017.10.037

Varghese, V., Magnani, L., Harada-Shoji, N., Mauri, F., Szydlo, R. M., Yao, S.,… Kenny, L. M. (2019). FOXM1 modulates 5-FU resistance in colorectal cancer through regulating TYMS expression. Sci Rep, 9(1), 1505. doi:10.1038/s41598-018-38017-0

Waldemer-Streyer, R. J., Reyes-Ordoñez, A., Kim, D., Zhang, R., Singh, N., & Chen, J. (2017). Cxcl14 depletion accelerates skeletal myogenesis by promoting cell cycle withdrawal. npj Regenerative Medicine, 2(1), 16017. doi:10.1038/npjregenmed.2016.17

Wang, Y., Liu, S., Yan, Y., Li, S., & Tong, H. (2019). SPARCL1 promotes C2C12 cell differentiation via BMP7-mediated BMP/TGF-β cell signaling pathway. Cell death & disease, 10(11), 852. doi:10.1038/s41419-019-2049-4

Wang, Z., Wang, X., Zou, H., Dai, Z., Feng, S., Zhang, M.,… Cheng, Q. (2020). The Basic Characteristics of the Pentraxin Family and Their Functions in Tumor Progression. Frontiers in Immunology, 11(1757). doi:10.3389/fimmu.2020.01757

Wicher, S. A., Prakash, Y. S., & Pabelick, C. M. (2019). Caveolae, caveolin-1 and lung diseases of aging. Expert Rev Respir Med, 13(3), 291–300. doi:10.1080/17476348.2019.1575733

Wolf, F. A., Angerer, P., & Theis, F. J. (2018). SCANPY: large-scale single-cell gene expression data analysis. Genome Biology, 19(1), 15. doi:10.1186/s13059-017-1382-0

Wolf, F. A., Hamey, F. K., Plass, M., Solana, J., Dahlin, J. S., Göttgens, B.,… Theis, F. J. (2019). PAGA: graph abstraction reconciles clustering with trajectory inference through a topology preserving map of single cells. Genome Biol, 20(1), 59. doi:10.1186/s13059-019-1663-x

Wu, S. A., Kersten, S., & Qi, L. (2021). Lipoprotein Lipase and Its Regulators: An Unfolding Story. Trends Endocrinol Metab, 32(1), 48–61. doi:10.1016/j.tem.2020.11.005

Xu, Q. Q., Qin, L. T., Liang, S. W., Chen, P., Gu, J. H., Huang, Z. G.,… Chen, J. B. (2020). The Expression and Potential Role of Tubulin Alpha 1b in Wilms’ Tumor. Biomed Res Int, 2020, 9809347. doi:10.1155/2020/9809347

Xu, X., Wilschut, K. J., Kouklis, G., Tian, H., Hesse, R., Garland, C.,… Pomerantz, J. H. (2015). Human Satellite Cell Transplantation and Regeneration from Diverse Skeletal Muscles. Stem Cell Reports, 5(3), 419–434. doi:10.1016/j.stemcr.2015.07.016

